# Picturing plant biodiversity from airborne environmental DNA

**DOI:** 10.1101/2024.01.11.571706

**Authors:** Anne-Céline Thuillet, Didier Morisot, Jean-François Renno, Nora Scarcelli, Julien Serret, Cédric Mariac

**Affiliations:** Institut de Recherche pour le Développement, UMR DIADE, 911 Avenue Agropolis, BP 64501, 34394 Montpellier Cedex 5, France; Jardin des Plantes de Montpellier, FACULTÉ DE MÉDECINE, 2 rue École de médecine, CS 59001, 34060 Montpellier Cedex 2, France

## Abstract

While eDNA approaches have gained interest over the past decades all types of organisms have not been addressed evenly. In particular terrestrial plants have been the subject of less attention. Here we address the possibility to represent plant biodiversity from airborne environmental DNA (eDNA) sampling and metabarcoding. We collected air using a biological air sampler in the Botanical Garden of Montpellier (France) and compared the list of revealed plant species to the botanical inventory of the Garden. Ninety-two plant species could be detected from three sampling points across the 4,6 ha of the Garden, after one hour sampling allowing to filter 9 m^3^ of air. We recorded the plants carrying flowers at the time of the experiment, which allowed us to estimate that plants flowering at the time of the sampling could be detected 10 times more easily than plants that were not, given the number of plants carrying flowers. However, flowering is far from being required as a vast majority of plants still was detected without flowering. We also show that not all species orders are detected with the same probability, tree species being better represented in the sample than herbal plants, given the number of trees present in the garden. Finally using diagnostic species, present only once in the garden, we estimate that the maximum sampling distance allowed by the biological air sampler is at least 110 m. Our study underlines that air sampling is a promising method for monitoring terrestrial plant biodiversity and highlights the parameters that should be adjusted to optimize the approach.

## INTRODUCTION

Current changing climate conditions accelerate the decline of biodiversity, all habitats and species confounded (Arneth et al, 2020). Our ability to efficiently monitor biodiversity changes is key in limiting negative impacts on ecosystems equilibrium (Hines and Pereira, 2021). However, traditional biodiversity surveys, based on species inventories, are time consuming, costly, and limited in terms of spatial scales as in their ability to detect rare species. In addition, the approach requires specialists to precisely identify taxa. It can also be particularly challenging in difficult or hostile environments, as water, or tropical forests (Zinger et al, 2020).

Over the last decades, sampling the environment instead of sampling organisms directly has proven to be an efficient approach to reveal local biodiversity (see Ruppert et al, 2019; Takahashi et al, 2023 for reviews). Based on the fact that all organisms release in their environment DNA (coming from debris, hair, feces, cells…), that can be collected, amplified and sequenced using metabarcoding, the approach has been applied for instance in the early detection of invasive species (Blackman et al, 2020), for the analysis and monitoring of ecological communities (Cilleros et al, 2019), or when direct sampling is difficult (Sigsgaard 2016). Since the rise of the method however, developments have unequally addressed the different types of organisms, leaving behind terrestrial plants (Banerjee et al, 2022; Johnson et al, 2023). Only about 4% of the studies based on eDNA sampling have been published since 2008 on contemporary vascular plants (representing about 160 studies, versus 4114 articles across all organisms, Banerjee et al, 2022). Among these, a handful of papers are based on air sampling to study pollen in relationship with human health (Folloni et al., 2012; Kraaijeveld et al., 2015; Korpelainen and Pietilainen 2017; Mohanty et al., 2017, Nuñez et al, 2017; Brennen et al, 2019; Banchi et al, 2020, Rowney et al, 2021), and a minority considered airborne eDNA as an approach to monitor terrestrial plant biodiversity (Johnson et al, 2019a, 2019b, 2021, Leontidou et al, 2021). Only two of these studies, to the best of our knowledge, really compare a listing of species revealed from airborne eDNA to a classic local botanical survey: one analyzes a short-grass rangeland (Johnson et al, 2021) and a second one focuses on a more complex alpine community, at a larger scale (Leontidou et al, 2021). It is thus needed to further explore the possibility of using airborne sampling to monitor terrestrial plant biodiversity, in order to better understand how this approach can be applied for ecological and conservation purposes.

The Botanical Garden of Montpellier is located in the city centre and was established in 1593. It covers 4,5ha and contains 5788 referenced specimens belonging to 3121 species of trees, shrubs, ornamental plants, naturalized plants, and weeds, that are distributed among 1120 Genera and 172 Families. Available information of this inventory covers the botanical name of each plant referenced (species, genus, family and sub-taxa), its vernacular name, its geographical origin, biological type (annual, perennial…), usage and observation, and its localization within the Garden. Localization is given by one or most often several parcel numbers for herbal plants (except for weeds that are not localised). Trees localization is on the contrary precisely known.

In this study, our objective was to provide a first picture of the way airborne eDNA manages to reveal plant biodiversity in a complex botanical context covering herbal plants, trees and bushes. To do so, we collected air samples in the Botanical Garden of Montpellier and analyzed these samples using a metabarcoding approach. We then compare the list of species as revealed by the obtained sequences to the list of plant species botanically described in the Garden. We also analyzed different parameters that may influence the method’s efficiency, by asking the following questions: (i) does sampling duration linearly increase the number of plants detected, given the used device, or is there an optimum? (ii) do we detect more species by applying two different DNA extraction’s protocols to separate heavy and light particles? (iii) are flowering plants at the time of the sampling more often detected than non-flowering plants (as it could be expected through the release of pollens)? (iv) are the different types of plants (herbal plants versus trees) evenly detected, given the number of each type present in the Garden? (v) at which maximum distance can we detect a plant species?

## RESULTS AND DISCUSSION

### Revealing plant species from air sampling

We sampled air in three locations of the Botanical Garden of Montpellier (Figure 1) using a biological air sampler (Coriolis ® microAir Sampler, Bertin Technologies). To determine the optimum volume of air to sample using this device, we analyzed three sampling durations (each being totalized over the three locations and repeated twice for duplication). We sampled three times 1 min, three times 3 min and three times 6 min, totalizing respectively 3 min, 9 min and 18 min of sampling in three tubes, and corresponding respectively to a total of 0,9 m^3^, 2,7 m^3^ and 5,4 m^3^ of air filtered. As we may have sampled pollen and debris but also cells or fragmented DNA, samples were centrifuged to separate heavy particles from more fragile material. DNA extractions were carried out on pellets on the one hand and on eluates on the other hand, as pollen DNA extraction requires a more aggressive protocol that might damage more fragile DNA. Extracted samples were analyzed using three barcodes: rbcL 230, rbcL260 and trnLg of respective lengths of 244 bp, 380 bp and 107 bp designed on two chloroplastic genes (rbcL and trnL) commonly used for plants and chosen for their representation in databases. Details and protocols are described in the Method section.

**Figure 1:**
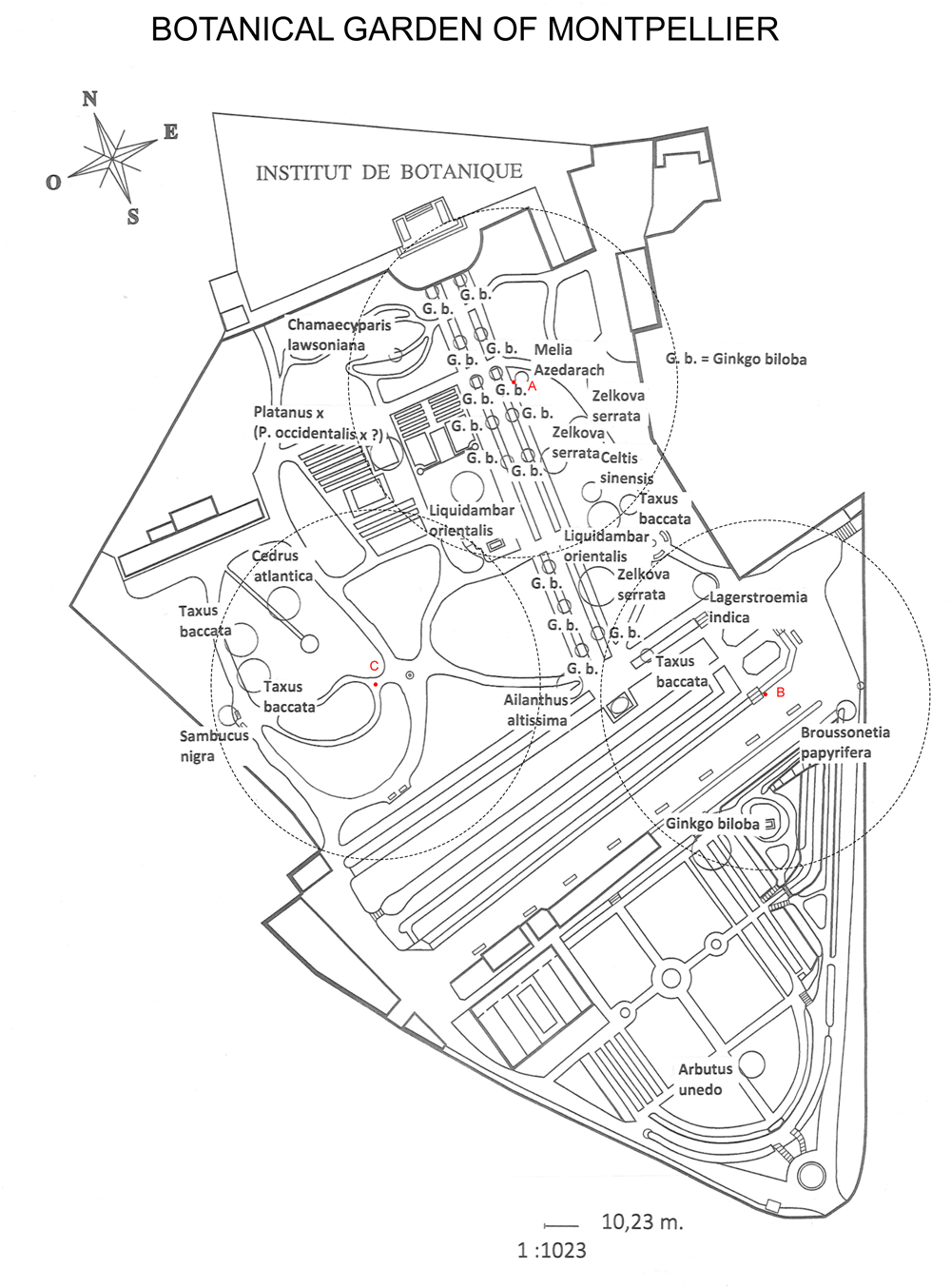
Tree species and sampling points localized on the map of the Botanical Garden. Trees are represented as plain circles. Circles size is representative of trees size. Red dots are the three sampling points (A, B, C). Dotted circles represent 50m radius areas around sampling points within which flowering was surveyed.

### Experimental factors affecting species detection

Figure 2 shows that the relationship between the sampling time and the number of species detected is not linear, but rather presents a significant optimum for 3 minutes of sampling per location (Wilcoxon test p<0.0001). This can be explained by the fact that collecting tubes that contain initially 15 ml of collecting buffer are agitated by a vortex that allows particles to fall into the tube. This vortex creates loss of liquid that should be compensated along the experiment. Thus, the longer sampling lasts, the more diluted the sample becomes, through the addition of buffer, required to maintain a 15 ml volume in the tube. Rarer species may not be sampled again after this buffer correction, resulting in a more difficult detection because of a higher dilution factor after a longer sampling time. A similar observation was made by Rufino de Sousa et al. (2020), using the same device for microbiological air sampling.

**Figure 2:**
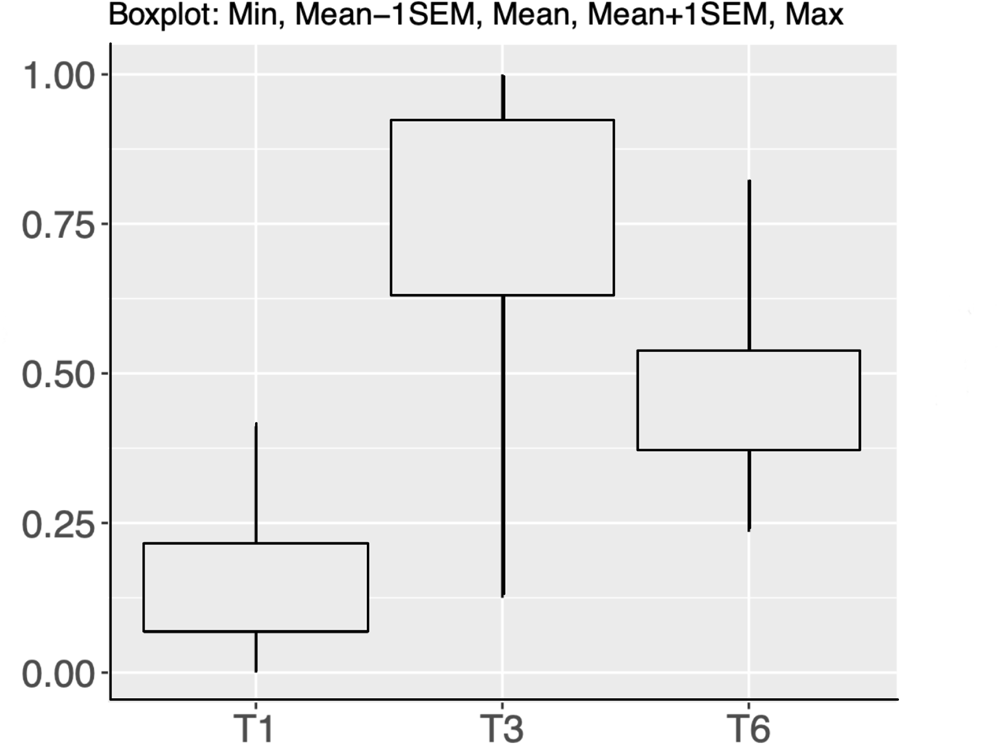
Boxplot showing the normalized number of species* (y axis) for all barcodes detected for each sampling duration (x axis). Duration are named after T1, T3 and T6 for 1min, 3min and 6 min per sampling location respectively. *In order to sum alpha diversity for the different barcodes, a Min-Max normalization was applied.

The comparison of both extraction types showed that almost all species identified from centrifugation eluates were also detected in pellets (16 among 20 species detected in eluates across all samples). Most species found in eluates were detected by the less resolutive barcode trnL. Four species were detected only in eluates, namely *Eleusine multiflora*, *Enneapogon lindleyanus*, *Phalaris truncata* and *Senna rugosa*, resp. 3 poaceae and a fabaceae. Only one of these species is described in the region of Montpellier (*Phalaris truncata*), and none is present in the Botanical Garden inventory. The identification of species may be subject to errors that will be discussed below. But given the low number of species that are specific to eluates and given the fact that they might even be misidentifications, we conclude from our results that DNA extracted from centrifugation eluate may be neglected from air sampling, and that, when applied to plant taxa, analyses may rely on a single DNA extraction protocol adapted to pellets. Following results only include pellet extractions output.

The three barcodes allowed the identification of different numbers of species, the average number of species identified being significantly higher with each of the rbcL barcodes than with the trnLg barcode (47 species for rbcL230 as for rbcL260 versus 21 species for trnLg, Wilcoxon test, p<0,01). This is consistent with the fact that the trnLg barcode being smaller, it is less resolutive in identifying species. We obtained little overlap between the species detected by the three barcodes, rather showing a good complementarity (Figure S1).

### Comparing biodiversity as revealed from eDNA to the botanical description of the Garden

We obtained 2,21 millions of filtered sequencing reads over all samples. Our sequencing effort is likely to be sufficient as supported by the rarefaction curves, reaching most of the time an asymptote at the maximum sequencing depth (Figure S2).

Despite only 60 minutes in total of collection sampling in three locations in the whole botanical garden, we identified a total of 92 species in our samples. Among these, 67 are present in the Botanical Garden (Table 1). The 25 species that are not found as present in the listings of the Botanical Garden comprises 5 species (namely *Trifolium repens* (clover)*, Torilis japonica* (Japanese hedge parsley), *Allium oleraceum* (garlic), *Alopecurus myosuroides* (black-grass) and *Populus tremula* (aspen)) that are common in France, and likely to exist in or near the Botanical Garden, according to the botanist of the Garden and to the species distributions data (french National Inventory of Natural Heritage (https://inpn.mnhn.fr)). Seventeen of the non-listed species are close to one or several other species belonging to the same genus and present in the Garden. Several hypotheses may explain this result. First, species may not be correctly labeled in databases, leading to misidentifications. Second, the species present in the garden do not have a barcode available, but a closely related species absent from the garden does, in which case the latter is returned as a query result when browsing databases. Finally, the sequences of some barcodes may not always be of full-length in the databases and the overlap might be insufficient to identify the right species. The three last species identified are more surprising. *Anacardium occidentale* is the cashew tree. It is only cultivated in tropical lands and no specimens exist in the Botanical Garden of Montpellier. However, the fruit of this species, the cashew nut, is widely consumed, and it is not impossible that we detected it. *Hura crepitans* is an amazonian species also not present in the Botanical Garden of Montpellier, but it has been found in another e-DNA study made in Peru by our lab. An isolated cross contamination is not to be excluded, although it was not found in our negative controls. *Acanthopsis spathularis* is a species native from South Africa and no explanation is available to support its detection: isolated foreign detection is nevertheless part of eDNA studies and may be imputed to imprecise assignation (Burian et al, 2021).

**Table 1:** Species identified in air samples.

| Species name | number<br>of reads<br>rbcL230 | number<br>of reads<br>rbcL260 | number<br>of reads<br>trnLg | total<br>number<br>of reads | flowering<br>(Y/N) | Type :<br>herbal<br>plant<br>(H)<br>Tree (T)<br>shrub/<br>bush (S) | number of<br>individuals<br>(for trees) |
| --- | --- | --- | --- | --- | --- | --- | --- |
| Acanthus mollis |  | 4 |  | 4 | N | H |  |
| Ailanthus altissimus | 12 | 2 | 1 | 15 | N | T | 1 |
| Ampelodesmos mauritanicus | 5 | 7 |  | 12 | N | H |  |
| Arbutus unedo |  |  | 12 | 12 | N | T | 1 |
| Arctotheca calendula |  | 11 |  | 11 | Y | H |  |
| Arenaria serpyllifolia |  | 17 |  | 17 | N | H |  |
| Bellis perennis | 7 | 107 |  | 114 | Y | H |  |
| Bromus erectus | 45 |  |  | 45 | N | H |  |
| Broussonetia papyrifera |  | 16618 | 106510 | 123128 | N | T | 1 |
| Carex pendula |  | 2 |  | 2 | Y | H |  |
| Cedrus atlantica |  | 95 |  | 95 | N | T | 1 |
| Celtis sinensis | 40374 |  |  | 40374 | N | T | 1 |
| Centranthus ruber |  |  | 44 | 44 | Y | H |  |
| Chamaecyparis lawsoniana | 248 |  |  | 248 | N | T | 1 |
| Chelidonium majus |  | 91 | 203 | 294 | Y | H |  |
| Cichorium intybus |  | 100 |  | 100 | N | H |  |
| Convolvulus arvensis |  | 7 |  | 7 | N | H |  |
| Coriaria myrtifolia | 363 |  | 13 | 376 | N | S |  |
| Crepis capillaris | 37 | 93 |  | 130 | N | H |  |
| Cynodon dactylon |  | 3 |  | 3 | N | H |  |
| Dactylis marina | 21 |  |  | 21 | N | H |  |
| Erodium cicutarium |  | 8 |  | 8 | N | H |  |
| Galium aparine |  | 3 |  | 3 | N | H |  |
| Ginkgo biloba | 6395 | 4100 |  | 10495 | Y | T | 15 |
| Helminthotheca echioides | 4 | 6 |  | 10 | N | H |  |
| Isatis tinctoria |  | 22 |  | 22 | Y | H |  |
| Lagerstroemia indica | 33 |  |  | 33 | N | T | 2 |
| Lamium purpureum | 5 |  |  | 5 | Y | H |  |
| Lapsana communis |  | 11 |  | 11 | N | H |  |
| Lathyrus cicera |  |  | 3 | 3 | N | H |  |
| Liquidambar orientalis |  | 197 |  | 197 | Y | T | 2 |
| Medicago arabica | 251 |  | 1 | 252 | N | H |  |
| Medicago constricta |  |  | 15 | 15 | N | H |  |
| Medicago lupulina | 696 | 112 |  | 808 | N | H |  |
| Medicago polymorpha |  | 53 |  | 53 | N | H |  |
| Medicago truncatula | 6 |  |  | 6 | N | H |  |
| Melia azedarach |  |  | 30 | 30 | N | T | 1 |
| Melilotus officinalis | 62 |  | 1268 | 1330 | N | H |  |
| Mercurialis annua | 17 | 10 | 16 | 43 | N | H |  |
| Nelumbo lutea |  | 7 |  | 7 | N | H |  |
| Papaver rhoeas | 17 |  |  | 17 | N | H |  |
| Parietaria judaica | 6786 |  |  | 6786 | N | H |  |
| Plantago lanceolata | 496 |  |  | 496 | N | H |  |
| Platanus occidentalis | 105628 |  |  | 105628 | N | T | 1 |
| Platanus racemosa |  |  | 1168 | 1168 | N | T |  |
| Poa annua | 692 | 472 |  | 1164 | N | H |  |
| Polycarpon tetraphyllum | 4 |  |  | 4 | N | H |  |
| Potentilla indica |  |  | 3 | 3 | N | H |  |
| Salpichroa organifolia |  | 11 |  | 11 | N | H |  |
| Sambucus nigra | 17 |  |  | 17 | N | S | 1 |
| Sanguisorba minor | 19 | 15 |  | 34 | N | H |  |
| Saponaria officinalis |  | 28 |  | 28 | N | H |  |
| Sherardia arvensis |  | 5 |  | 5 | N | H |  |
| Solanum tuberosum |  | 20 |  | 20 | N | H |  |
| Stellaria media | 179 | 136 |  | 315 | N | H |  |
| Taxus baccata |  | 718 |  | 718 | N | T | 4 |
| Torilis nodosa |  | 7 |  | 7 | N | H |  |
| Trifolium scabrum |  | 4 | 2474 | 2478 | N | H |  |
| Urtica membranacea |  | 9 |  | 9 | N | H |  |
| Valerianella locusta | 9 | 4 |  | 13 | N | H |  |
| Veronica arvensis | 140 | 62 | 1 | 203 | N | H |  |
| Veronica persica | 23 |  |  | 23 | N | H |  |
| Veronica sublobata |  | 34 |  | 34 | N | H |  |
| Viburnum tinus | 58 |  |  | 58 | Y | S |  |
| Xanthoceras sorbifolium |  |  | 11 | 11 | N | S |  |
| Zea mays | 14 |  |  | 14 | N | H |  |
| Zelkova serrata | 1413 | 10 |  | 1423 | N | T | 3 |

The 67 species detected among 1585 species present in the Botanical Garden and identifiable because they have at least one available barcode represent a 4% rate of detection given our sampling effort. We used one hour to filter 18 m^3^ of air, spanning 3 locations in the Botanical Garden of Montpellier (4,6 ha). From these data, it is possible to estimate a theoretical number of 22 species detected per location, about 4 species per m^3^, or still about 15 species per ha. In short, at least a 24 fold-increased sampling effort would have been required to detect all the botanical garden’s species.

### Ecological factors affecting species detection

Beyond the duration of the sampling, the volume of air filtered, or the number of locations explored, the ecology of the targeted species also is likely to affect their detection, as well as their physiology, repartition, surface, density and biomass. For plants, one may hypothesize that anemophilous pollination could increase detectability, although it has been shown that non pollinating or entomophilous species at the time of the sampling are also detectable through air sampling (Johnson et al, 2019b). More generally, biomass generally has an impact, bigger specimens or denser populations releasing more DNA than smaller individuals or sparser populations. Finally, a specimen’s detectability via eDNA is also likely to be influenced by its distance to the sampling location, the more remote, the less detectable.

At the time of the sampling 98 species were at flowering stage within a 50 meters radius around each of our three sampling points. Among these 98 flowering plants 62 are represented in databases by at least one barcode. Among the 67 specimens identified at species level in our eDNA sampling, and confirmed to be present in the Botanical Garden, 10 were flowering at the time of the sampling. We thus detected 10 flowering species among 62 detectable species, i.e. a proportion of 0,16. A z-test describes that this proportion is significantly higher than expected if flowering had no impact on species detectability (67/1598=0,04, z=4,6894, p<0,00001, two-tailed test). It should be noted that the survey of flowering species does not cover the totality of the garden, but only about half the surface. As a consequence, the proportion of detected flowering plants could be overestimated (if additional detectable species are flowering outside of the surveyed area). However, if we hypothesize that by doubling the surface, as many flowering species have been missed by the survey, the z-test would still be significant (p<0,02), which makes our conclusion rather conservative.

A comparable analysis can be led regarding the type of species that can be detected. Specifically, it is possible to compare the number of big specimens (i.e. trees), versus smaller ones, the former being expected to release more DNA in the air. We observe that 14 trees were detected among 141 detectable ones (present in the Garden and represented in databases by at least one barcode). This proportion of 0,09 trees is significantly higher than expected if size had no impact on detectability (67/1598=0,04, z=3,3986, p<0,0007, two-tailed test).

Detectability may also be analyzed through the number of reads found in the samples. As this variable is not normally distributed in our sample, we applied the non-parametric test of Sheirer Ray-Hare to analyze the effect of flowering and the effect of specimen size on the number of reads, following the same principle as an analysis of variance. Regarding specimen size, we considered two levels: herbal plants and trees. Shrubs were classified under “herbal plants” so that only big trees are considered as plants with a significantly higher biomass per individual. Biomass was found to have a significant effect on the number of reads (p=0,0006), while neither flowering nor the interaction biomass x flowering had any effect (p=0,82 and p=0,68 respectively). The results of this analysis holds (p=0,01 for plant type, and NS for flowering and interaction)) when discarding two trees presenting extreme values of read numbers (*Broussonetia papyrifera*, 123128 reads; *Platanus occidentalis*, 105608 reads), and one tree species known to be represented by 18 individuals in the garden (*Ginko biloba*), while other trees are represented by 1 to 4 individuals.

Finally, we considered testing the effect of distance on the number of reads, by using the 10 trees that were detected in our eDNA samples, and that are represented by one individual precisely localized in the garden. We found negative correlations between the number of reads in the samples and, on the one hand, the mean distance between trees and sampling points and with, on the other hand, the distance between trees and their closest sampling point, but none was significant (see supplementary Figure S3), probably because of a lack of power, too many confounding factors, and the fact that our samples are pooled over the three sampling points. However, from our data, we may report that the more remote tree from our sampling points is located 110 m away from the closest sampling point to it. This represents the maximum measurable distance within which we could detect plants.

### Conclusions

While eDNA approaches are seen as very promising for biodiversity monitoring, applications in plants are still scarce in the literature (Banerjee et al, 2022; Johnson et al, 2023). With our study, we confirm that the potential of eDNA for plant studies is as promising as for other organisms, based on the detection of 92 species from a relatively small sampling effort, spanning only 3 locations and including 6 samples for a total of 18m^3^ of air analyzed. However, parameters should be studied to optimize the use of eDNA approaches based on air sampling for plants: the ecology of plants appears to impact their detectability, through their biomass or their phenological status. Stratified sampling in space and time or roving sampling could be carried out to measure the effect of these different parameters on the effectiveness of the plant biodiversity census through air sampling. Finally, although the detection distance can be quite far (minimum 110 m), a tighter meshing should increase the chances of species detection, as long as the sampling effort is still advantageous compared with a conventional biodiversity inventory.

## METHODS

### Flowering characterization

We surveyed species that were flowering at the time of the experiment within a 50-meter radius of each of our three sampling points (see Figure 1). A total of 99 species were shedding pollen, among which 79 are entomophilous, 19 are anemophilous species, and one has a mixed reproductive system.

### Sample collection

Air was collected at the Botanical Garden of Montpellier (France, 43° 36.853’N 3° 52.319’E) on April, 15th 2022. We used a biological air sampler (Coriolis ® micro Air Sampler, Bertin Technologies) in three locations of the garden positioned 100 meters apart from each other (see Figure 1). The device was positioned at a height of 120 cm above the ground. It vacuums air over a sterile cone containing 15 ml of sterile liquid collection (Bertin Technologies), by creating a vortex that traps airborne particles into the cone. In order to evaluate the effect of the volume of air sampled on the number of species detected, we applied three sampling durations: 1mn, 3mn, and 6mn per location, i.e. 3mn, 9mn and 18mn of sampling duration pooled over the three locations in each of the three cones, and corresponding respectively to a total of 0,9 m^3^, 2,7 m^3^ and 5,4 m^3^ of air filtered, the device vacuuming 300 liters of air per minute. A replicate was done for each sampling, resulting in 6 samples in total. In each location, a refill was made up to 15ml before air collection if necessary. A negative field control added as a cone filled with 15 ml of liquid collection in the field, and opened at each collecting point, i.e. 6 times. After each air collection the cones were stored at 4 degrees. At the end of sampling, the cones were transported to the laboratory and stored at −20°C until DNA extraction.

### DNA extraction

For each of the 7 cones, the 15 ml of liquid collection were centrifuged at 8700 rcf for 20 mins. The supernatant was transferred to a new 50 ml tube. Two extractions per sample were performed, one on the pellet and one on the collected supernatant. The pellet was resuspended in 400 µl of 1X longmire buffer (Longmire et al., 1997), then transferred to a 1.5 ml tube containing 200 µl of ceramic beads for sonication with a bioruptor (2 cycles of 30 s on, 30 s off). Then 15 µl of Proteinase K (20 mg/ml) and 80 µl SDS 10% were added, and lysis performed overnight at 56°C on an eppendorf thermomixer with an agitation at 500 rpm. Purification was conducted by adding 600 µl chloroform isoamyl alcohol (24:1) and, after mixing, centrifuged at 10000 rcf for 10 mins. Supernatants were carefully collected and precipitated with 500 µl isopropanol and 60 µl 3M Na acetate pH5 followed by a centrifugation at 8700 rcf for 30 mins at 4°C. The pellets were rinse with 70% ethanol and centrifuged at 12000 rcf for 30 mins. After supernatants elimination, pellets were air dried and resuspended with 30µl ultra-pure water.

The supernatant was treated in the same way but without bioruptor lysis and with the addition of 3 ml 5X longmire buffer, 15 µl Proteinase K and 0.3 g SDS. Centrifugations were performed at 8700 rcf for purification with chloroform isoamyl alcohol, precipitation with isopropanol and rinsing with 70% ethanol.

### PCR amplification and sequencing

Metabarcoding was performed with three different chloroplastic barcodes: trnLg of 107 bp (Taberlet et al., 2007) (F: 5’-GGGCAATCCTGAGCCAA-3’; R: 5’- CCATTGAGTCTCTGCACCTATC-3’), rbcL230 of 244 bp (F: 5’- ATGTCACCACAAACAGAGACTAAAGC-3’ ; R: 5’- CCTTTGTAACGATCAAGRCTGGTAAG-3’) and rbcL260 of 380 bp (ref F: 5’- CTTACCAGYCTTGATCGTTACAAAGG-3’; R: 5’-GTAAAATCAAGTCCACCRCG-3’). The primers used for amplification of the two rbcL locus were taken from (García-Robledo et al., 2013) and are consecutive on the rbcL loci meaning that the reverse primer of rbcL230 is the reverse complement of the forward primer of rbcL260.

Library preparations were performed at ADNid (Montferrier-sur-Lez, France; http://www.adnid.fr) according to the two-step PCR described by Galan et al. 2018 (see STable 1 for details). The first PCR were performed using Type-it Microsatellite PCR Kit (Qiagen 206243) and following the manufacturer’s instructions in a final volume of 15 µl and with 2 µl of eDNA. The amplification PCR for the rbcL makers was performed under the following conditions: preincubation at 95°C for 5 min; 40 cycles (30 sec at 95°C, 1 min at 50°C, 30 sec at 72°C); final extension 30 min at 60°C. The amplification PCR for the trnLg maker were performed under the following conditions: preincubation at 95°C for 5 min; 35 cycles (30 sec at 95°C, 1 min at 55°C, 30 sec at 72°C); final extension 30 min at 60°C.

The three barcodes were amplified separately with an extension added on each forward primer (5’-TCGTCGGCAGCGTCAGATGTGTATAAGAGACAG-3’) and reverse primer (5’-GTCTCGTGGGCTCGGAGATGTGTATAAGAGACAG-3’). In order to link the sequencing adaptors and double indexed libraries the second PCR step were performed using TG Nextera® XT Index Kit (FC-131-200(1,2,3) and the Kappa Hifi Hot Start (Roche KK2602) following the manufacturer’s instructions. The second PCR amplification was performed under the following conditions: preincubation 3min at 95°C ; 12 cycles (30 sec at 95°C, 30 sec at 55°C, 30 sec at 72°C) ; final extension 5 min at 72°C.

In addition to the negative field controls (E-TNC and C-TNC, resp. for eluate and pellet extractions of the negative control cone), two extraction negative controls (E-TNE and C-TNE, made on 15 ml of fresh buffer and extracted at the same time as the samples) and two PCR negative controls (ultrapure water instead of DNA) were amplified on each of the three barcodes. Each barcode were equimolary pooled, then purified and paired end sequenced (250 bp) using Miseq Illumina platform at the ARCAD (Agropolis Resource Center for Crop Conservation, Adaptation and Diversity, https://www.arcad-project.org) facilities.

### Bioinformatics methods and data filtering

Amplicon sequences were analyzed using the FROGS pipeline version 3.1 (Escudié et al., 2018) with standard operating procedures for data cleaning (removing of reads without primers and chimeric sequences). For each library the three barcodes were split according to their primer sequences and size (see STable 1 for details). Clustering into operational taxonomic units (OTUs) was performed with Swarm (Mahé et al., 2014) using an aggregation distance of 1. After chimera removal, OTUs were kept if present in at least one library and with a minimum abundance of three, two and one sequence for rbcL230, rbcL260 and trnLg, respectively. In addition, we applied a very stringent assignment protocol, setting identity to 100%.

Taxonomic assignation of OTUs was performed by blastn (version2.8.1) against rbcL and trnL reference databases. The first one (rbcL) was public (Bell et al., 2017) and contains 38464 entries representing 26034 different plant species worldwide distributed. The second one (trnL) was built with sequences mined from GenBank and Bold contains 169989 entries representing 61368 plant species world wild distributed.

Among the 3121 species currently recorded in the JDP, 845 have both barcodes, 1391 and 1084 are represented only in the trnL and rbcL databases, respectively. Overall, 1598 species present in the botanical garden have at least one barcode available in one of these two reference databases.

Results from Blastn were parsed using the MEGAN software (Huson et al., 2016) version 6.24.11 using the naïve LCA algorithm and parameters showed in STable 2.

Negative controls (field, extraction and PCR) were used to identify and remove contaminating reads in sample libraries with the R package Microdecon (McKnight et al., 2019).

In order to assess the relevance of our sequencing effort, rarefaction curves were performed for each library using the R Vegan 2.5.6 package (Oksanen et al., 2016).

### Statistical analyses

Wilcoxon test and z test were performed using the R package Rstatix 0.7.2

Z-test was calculated as

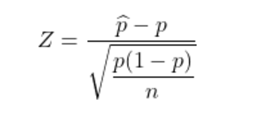

Where ^p is the proportion of the tested sample, p is the hypothesized proportion, and n is the sample size.

The Scheirer-Ray-Hare test was performed in R following Holmes et al (2016).

## Supporting information

supplementary_figS1_toS3_Stable1

## MATERIALS AVAILABILITY

This study did not generate new unique reagents.

## DATA AND CODE AVAILABILITY

Sequencing reads were deposited in the National Center for Biotechnology Information Sequence Read Archive (BioProject ID: PRJNA1033821, BioSample accessions: SAMN38046846-SAMN38046868). This paper does not report original code. Any additional information required to reanalyze the data reported in this paper is available from the lead contacts upon request.

## AUTHORS CONTRIBUTION

C. M., A.-C. T., J.-F. R. and N. S. designed the experiment; all of the authors performed field work; C.M., N. S. and A.-C. T performed the laboratory work; C. M. and A.-C. T. performed the analysis; D. M. facilitated the field work and performed the botanical identifications. All of the authors contributed to manuscript preparation.

## AKNOWLEDGMENTS

We thank the Botanical Garden of Montpellier for authorizing the sampling and for providing information regarding the specimens of the plants conserved in the Garden. This work was funded by the Agropolis foundation (2101-051).

## REFERENCES

1. Arneth A., Y.-J. Shin, P. Leadley, C. Rondinini, E. Bukvareva, et al., 2020 Post-2020 biodiversity targets need to embrace climate change. Proceedings of the National Academy of Sciences 117: 30882–30891. 10.1073/pnas.2009584117

2. Banchi E., C. G. Ametrano, E. Tordoni, D. Stanković, S. Ongaro, et al., 2020 Environmental DNA assessment of airborne plant and fungal seasonal diversity. Science of The Total Environment 738: 140249. 10.1016/j.scitotenv.2020.140249

3. Bell K. L., V. M. Loeffler, and B. J. Brosi, 2017 An rbcL reference library to aid in the identification of plant species mixtures by DNA metabarcoding1. Appl Plant Sci 5: apps.1600110. 10.3732/apps.1600110

4. Blackman R. C., K. K. S. Ling, L. R. Harper, P. Shum, B. Hänfling, et al., 2020 Targeted and passive environmental DNA approaches outperform established methods for detection of quagga mussels, *Dreissena rostriformis bugensis* in flowing water. Ecol. Evol. 10: 13248– 13259. 10.1002/ece3.6921

5. Brennan G. L., C. Potter, N. de Vere, G. W. Griffith, C. A. Skjøth, et al., 2019 Temperate airborne grass pollen defined by spatio-temporal shifts in community composition. Nat Ecol Evol 3: 750–754. 10.1038/s41559-019-0849-7

6. Burian A., Q. Mauvisseau, M. Bulling, S. Domisch, S. Qian, et al., 2021 Improving the reliability of eDNA data interpretation. Molecular Ecology Resources 21: 1422–1433. 10.1111/1755-0998.13367

7. Cilleros K., A. Valentini, L. Allard, T. Dejean, R. Etienne, et al., 2019 Unlocking biodiversity and conservation studies in high-diversity environments using environmental DNA (eDNA): A test with Guianese freshwater fishes. Mol Ecol Resour 19: 27–46. 10.1111/1755-0998.12900

8. Escudié F., L. Auer, M. Bernard, M. Mariadassou, L. Cauquil, et al., 2018 FROGS: Find, Rapidly, OTUs with Galaxy Solution, (B. Berger, Ed.). Bioinformatics 34: 1287–1294. 10.1093/bioinformatics/btx791

9. Folloni S., D.-M. Kagkli, B. Rajcevic, N. C. C. Guimarães, B. Van Droogenbroeck, et al., 2012 Detection of airborne genetically modified maize pollen by real-time PCR. Molecular Ecology Resources 12: 810–821. 10.1111/j.1755-0998.2012.03168.x

10. Galan M., J.-B. Pons, O. Tournayre, É. Pierre, M. Leuchtmann, et al., 2018 Metabarcoding for the parallel identification of several hundred predators and their prey: Application to bat species diet analysis. Molecular Ecology Resources 18: 474–489. 10.1111/1755-0998.12749

11. García-Robledo C., D. L. Erickson, C. L. Staines, T. L. Erwin, and W. J. Kress, 2013 Tropical Plant–Herbivore Networks: Reconstructing Species Interactions Using DNA Barcodes. PLoS ONE 8.

12. Hines J., and H. M. Pereira, 2021 Biodiversity: Monitoring trends and implications for ecosystem functioning. Current Biology 31: R1390–R1392.

13. Holmes, D., Moody, P., Dine, D., & Trueman, L. (2016). *Research Methods for the Biosciences* (Third edition). Oxford University Press. 488pp. ISBN: 9780198728498

14. Huson D. H., S. Beier, I. Flade, A. Górska, M. El-Hadidi, et al., 2016 MEGAN Community Edition - Interactive Exploration and Analysis of Large-Scale Microbiome Sequencing Data, (T. Poisot, Ed.). PLOS Computational Biology 12: e1004957. 10.1371/journal.pcbi.1004957

15. Johnson M. D., R. D. Cox, and M. A. Barnes, 2019a Analyzing airborne environmental DNA: A comparison of extraction methods, primer type, and trap type on the ability to detect airborne eDNA from terrestrial plant communities. Environmental DNA 1: 176–185. 10.1002/edn3.19

16. Johnson M. D., R. D. Cox, and M. A. Barnes, 2019b The detection of a non-anemophilous plant species using airborne eDNA, (H. Doi, Ed.). PLoS ONE 14: e0225262. 10.1371/journal.pone.0225262

17. Johnson M. D., R. D. Cox, B. A. Grisham, D. Lucia, and M. A. Barnes, 2021 Airborne eDNA Reflects Human Activity and Seasonal Changes on a Landscape Scale. Front. Environ. Sci. 8. 10.3389/fenvs.2020.563431

18. Johnson M. D., A. D. Katz, M. A. Davis, S. Tetzlaff, D. Edlund, et al., 2023 Environmental DNA metabarcoding from flowers reveals arthropod pollinators, plant pests, parasites, and potential predator–prey interactions while revealing more arthropod diversity than camera traps. Environmental DNA 5: 551–569. 10.1002/edn3.411

19. Korpelainen H., and M. Pietiläinen, 2017 Biodiversity of pollen in indoor air samples as revealed by DNA metabarcoding. Nordic Journal of Botany 35: 602–608. 10.1111/njb.01623

20. Kraaijeveld K., L. A. D. Weger, M. V. Garc, H. Buermans, J. Frank, et al., 2015 Efficient and sensitive identification and quantification of airborne pollen using next-generation DNA sequencing. 9.

21. Leontidou K., D. Vokou, A. Sandionigi, A. Bruno, M. Lazarina, et al., 2021 Plant biodiversity assessment through pollen DNA metabarcoding in Natura 2000 habitats (Italian Alps). Sci Rep 11: 18226. 10.1038/s41598-021-97619-3

22. Longmire J. L., M. Maltbie, and R. J. Baker, 1997 Use of “Lysis Buffer” in DNA isolation and its implication for museum collections/

23. Mahé F., T. Rognes, C. Quince, C. De Vargas, and M. Dunthorn, 2014 Swarm: robust and fast clustering method for amplicon-based studies. PeerJ 2: e593. 10.7717/peerj.593

24. McKnight D. T., R. Huerlimann, D. S. Bower, L. Schwarzkopf, R. A. Alford, et al., 2019 microDecon: A highly accurate read-subtraction tool for the post-sequencing removal of contamination in metabarcoding studies. Environmental DNA 1: 14–25. 10.1002/edn3.11

25. Mohanty R. P., M. A. Buchheim, J. Anderson, and E. Levetin, 2017 Molecular analysis confirms the long-distance transport of Juniperus ashei pollen. PLOS ONE 12: 1–13. 10.1371/journal.pone.0173465

26. Núñez A., G. Amo de Paz, Z. Ferencova, A. Rastrojo, R. Guantes, et al., 2017 Validation of the Hirst-Type Spore Trap for Simultaneous Monitoring of Prokaryotic and Eukaryotic Biodiversities in Urban Air Samples by Next-Generation Sequencing, (A. J. McBain, Ed.). Appl Environ Microbiol 83. 10.1128/AEM.00472-17

27. Oksanen J., F. G. Blanchet, M. Friendly, R. Kindt, P. Legendre, et al., 2016 *vegan: Community Ecology Package. Ordination methods, diversity analysis and other functions for community and vegetation ecologists*. The Comprehensive R Archive Network, California, Berkeley, USA.

28. Rowney F. M., G. L. Brennan, C. A. Skjøth, G. W. Griffith, R. N. McInnes, et al., 2021 Environmental DNA reveals links between abundance and composition of airborne grass pollen and respiratory health. Current Biology 31: 1995–2003.e4. 10.1016/j.cub.2021.02.019

29. Ruppert K. M., R. J. Kline, and M. S. Rahman, 2019 Past, present, and future perspectives of environmental DNA (eDNA) metabarcoding: A systematic review in methods, monitoring, and applications of global eDNA. Global Ecology and Conservation 17: e00547. 10.1016/j.gecco.2019.e00547

30. Sigsgaard E. E., I. B. Nielsen, S. S. Bach, E. D. Lorenzen, D. P. Robinson, et al., 2016 Population characteristics of a large whale shark aggregation inferred from seawater environmental DNA. Nature Ecology & Evolution 1: 0004. 10.1038/s41559-016-0004

31. Sousa N. R. de, L. Shen, D. Silcott, C. J. Call, and A. G. Rothfuchs, 2020 Operative and Technical Modifications to the Coriolis® µ Air Sampler That Improve Sample Recovery and Biosafety During Microbiological Air Sampling. Annals of Work Exposures and Health 64: 852–865.

32. Taberlet P., E. Coissac, F. Pompanon, L. Gielly, C. Miquel, et al., 2006 Power and limitations of the chloroplast trn L (UAA) intron for plant DNA barcoding. Nucleic Acids Research 35: e14–e14. 10.1093/nar/gkl938

33. Takahashi M., M. Saccò, J. H. Kestel, G. Nester, M. A. Campbell, et al., 2023 Aquatic environmental DNA: A review of the macro-organismal biomonitoring revolution. Science of The Total Environment 873: 162322. 10.1016/j.scitotenv.2023.162322

34. Zinger L., J. Donald, S. Brosse, M. A. Gonzalez, A. Iribar, et al., 2020 Chapter Nine - Advances and prospects of environmental DNA in neotropical rainforests, pp. 331–373 in Tropical Ecosystems in the 21st Century, Advances in Ecolxogical Research. edited by Dumbrell A. J., Turner E. C., Fayle T. M. Academic Press.

