## supplementary_figS1_toS3_Stable1 for "Picturing plant biodiversity from airborne environmental DNA"

SUPPLEMENTARY FIGURES LEGENDS

**Figure S1:** Venn diagram showing the number of species detected by each barcode and by two or three barcodes.

**Figure S2:** Rarefaction curves for from left to right trnL, rbcL230 and rbcL260 reads. Taxa richness is represented on the y axis. Sample size is represented on the x axis. Each curve corresponds to a sample: C codes for pellets, E for eluates; 1, 3 and 6 code for sampling durations of 1 min, 3 min and 6 min per location; a and b code for duplications. Mean.blank code for control samples treated together.

**Figure S3:** Plot A represents for ten trees present once in the Garden and detected in the eDNA samples, the relationship between the number of reads (y) and the mean distance between trees and the three sampling points (x). Plot B represents the same relationship considering the distance between trees and their closest sampling point (xmin).

Figure S1

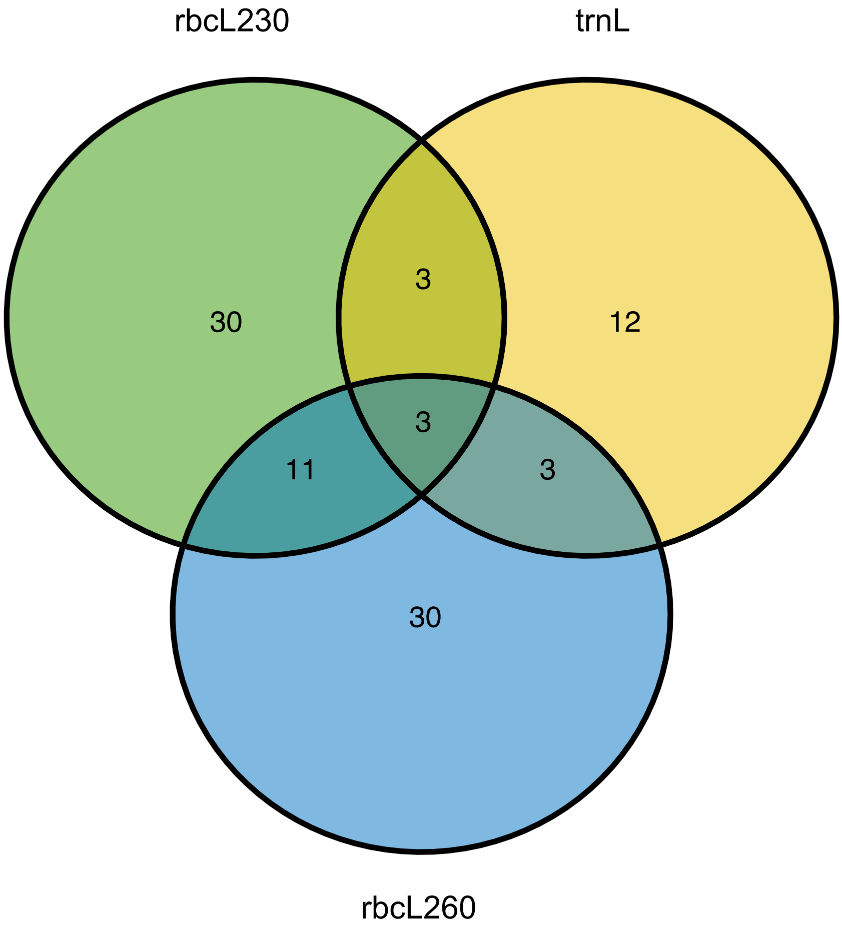

Figure S2

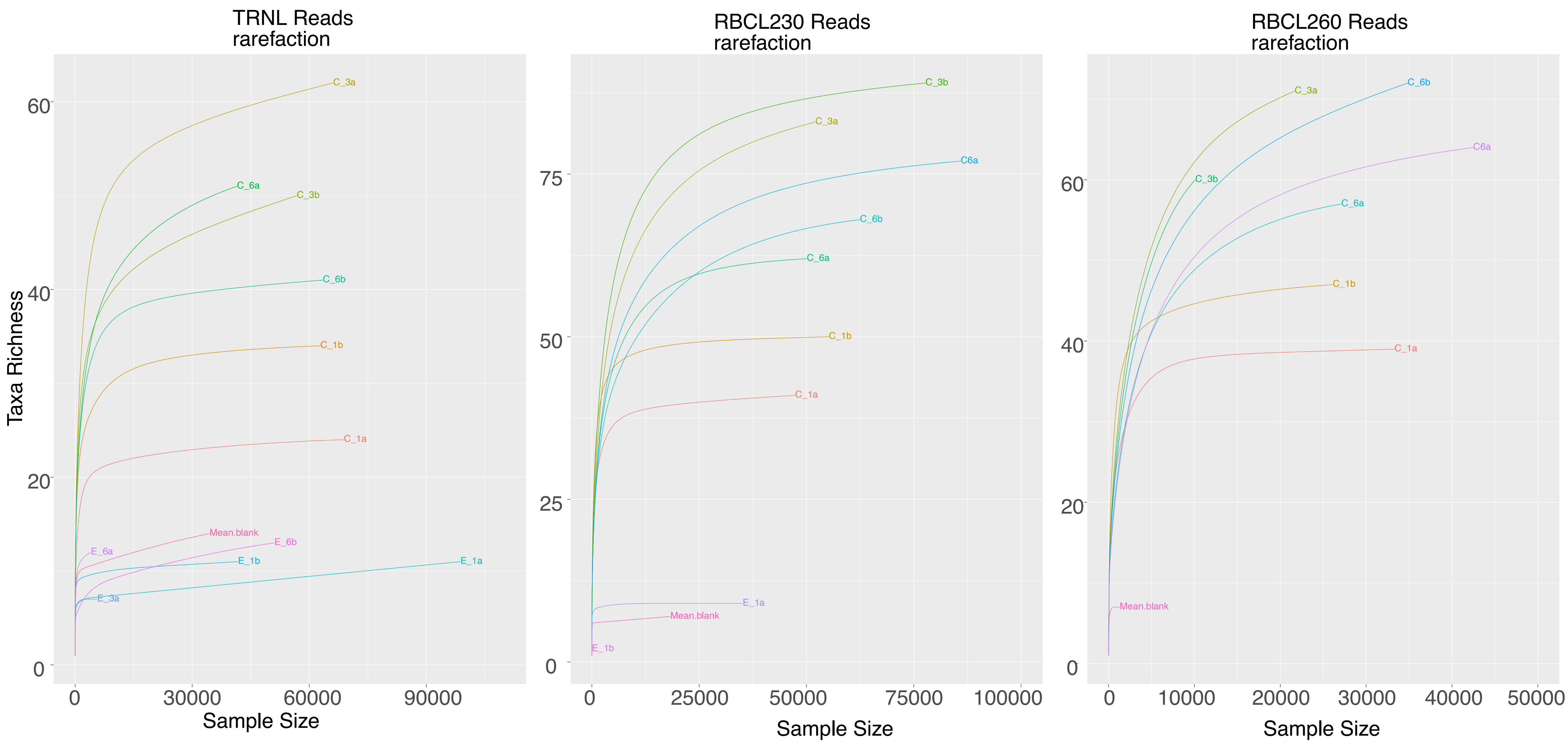
TRNL Reads
rarefaction
Taxa Richness
Sample Size
0
20
40
60
0
30000
60000
90000
C_3a
C_6a
C_3b
C_6b
C_1b
C_1a
Mean.blank
E_6b
E_6a
E_1b
E_1a
E_3a
RBCL230 Reads
rarefaction
0
25
50
75
0
25000
50000
75000
100000
C_3b
C_3a
C6a
C_6b
C_6a
C_1b
C_1a
E_1a
Mean.blank
E_1b
RBCL260 Reads
rarefaction
0
20
40
60
0
10000
20000
30000
40000
50000
C_3a
C_6b
C6a
C_3b
C_6a
C_1b
C_1a
Mean.blank

Figure S3

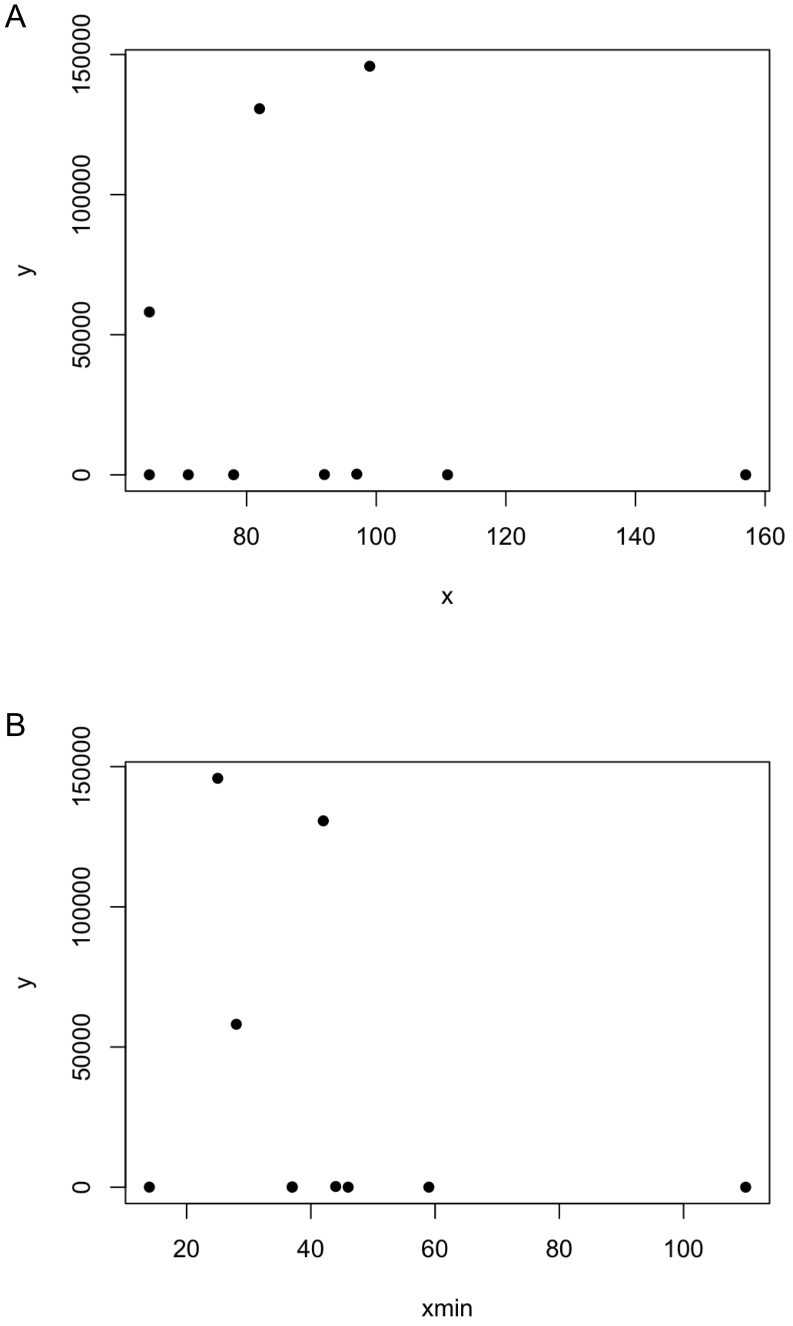
A
150000
100000
50000
0
y
80
100
120
140
160
x
B
150000
100000
50000
0
y
20
40
60
80
100
xmin

STable 1 : PCR primers, amplification conditions and Frogs parameters used for each of the three barcodes.

| | |
|---|---|
| | Forward Primer (red) with extension (Black) |
| rbcL230 | TCGTCGGCAGCGTCAGATGTGTATAAGAGACAGATGTCACCACAAACAGAGACTAAAGC |
| rbcL260 | TCGTCGGCAGCGTCAGATGTGTATAAGAGACAGCTTACCAGYCTTGATCGTTACAAAGG |
| TrnLgh (Sper01) | TCGTCGGCAGCGTCAGATGTGTATAAGAGACAGGGGCAATCCTGAGCCAA |
| | Reverse Primer (red) with extension (Black) |
| rbcL230 | GTCTCGTGGGCTCGGAGATGTGTATAAGAGACAGCCTTTGTAACGATCAAGRCTGGTAAG |
| rbcL260 | GTCTCGTGGGCTCGGAGATGTGTATAAGAGACAGGTAAAATCAAGTCCACCRCG |
| TrnLgh (Sper01) | GTCTCGTGGGCTCGGAGATGTGTATAAGAGACAGCCATTGAGTCTCTGCACCTATC |
| | Amplicon length |
| rbcL230 | 244 |
| rbcL260 | 380 |
| TrnLgh(Sper01) | 107 |
| | Tm |
| rbcL230 | 52 |
| rbcL260 | 52 |
| TrnLgh(Sper01) | 52 |
| | Number of cycles PCR1 |
| rbcL230 | 40 |
| rbcL260 | 40 |
| TrnLgh(Sper01) | 40 |
| | FROGS parameters (minAmpliconSize,maxAmpliconSize,fivePrimPrimer,threePrimPrimer,Read1 size,Read2 size,expected AmpliconSize) |
| rbcL230 | 200, 280, ATGTCACCACAAACAGAGACTAAAGC,CTTACCAGYCTTGATCGTTACAAAGG,250,250, 244 |
| rbcL260 | 300, 480, CTTACCAGYCTTGATCGTTACAAAGG,CGYGGTGGACTTGATTTTAC,250,250, 380 |
| TrnLgh (Sper01) | 80, 200, GGGCAATCCTGAGCCAA,GATAGGTGCAGAGACTCAATGG,150,150, 110 |

STable 2 : Megan parameters used for blastn parsing of the three barcodes

| | Min read number | Min score | Max Expected | Min Percent Identity (%) | Top Percent (%) | Min Support |
|---|---|---|---|---|---|---|
| trnL | 1 | 80 | 0,01 | 100 | 95 | 1 |
| rbcL230 | 3 | 200 | 0,01 | 100 | 100 | 1 |
| rbcL260 | 2 | 300 | 0,01 | 100 | 100 | 1 |
